# Molecular epidemiology of *Candida africana* isolates collected from vagina swabs in French Guiana

**DOI:** 10.1101/2023.12.13.571500

**Authors:** J. Bigot, Y. Kalboussi, Y. Bonkoto Nkoy, A. Benmostefa, S. Vellaissamy, L. Benzerara, V. Sainte-Rose, D. Blanchet, M. Demar, J. Guitard, C. Hennequin

## Abstract

*Candida africana* corresponds to the clade 13 of *Candia albicans*. It has been mostly involved in vulvovaginal candidiasis worldwide but few data exist in South America. The aim of our study was to investigate the prevalence of *C. africana* in women living in French Guiana. For this, we first set up a fluorescent-intercalating-dye-real time PCR targeting the hyphal wall protein 1 gene. The test was applied to 212 *C. albicans* isolates collected from May to August 2019 from vaginal swabs, allowing the identification of 6 women harboring *C. africana* (8 isolates). No demographics or clinical-specific features could be demonstrated. Genetic diversity of those isolates was analyzed through multilocus sequence typing and showed that diploid sequence type 182 was predominant (n=6) and allowed the report of a new diploid sequence type.

## Introduction

*Candida albicans*, an endogenous yeast part of the digestive and vaginal microbiota, is the predominant cause of vulvovaginal candidiasis (VVC), an infection affecting an estimated 138 million women annually (Denning et al. 2018). In 1995, atypical germ‐tube positive *C. albicans* isolates, recovered from African patients suffering VVC, led to the establishment of a new species called *Candida africana* (Tietz et al. 1995). Thereafter molecular typing demonstrated that *C. africana* does not meet the criteria for being individualized as a distinct species but in fact corresponds to the clade 13 of the *C. albicans* species (Sharma et al. 2014; Borman et al. 2013; Felice et al. 2016). For an easier reading of this article, strains belonging to the clade 13 of *C. albicans* will be named *C. africana* strains while strains from the other clades will be named *C. albicans*. Most of the previous surveys point out the isolation of *C. africana* from vaginal samples, while isolation from other body sites is rare (Gharehbolagh et al. 2020). *C. africana* have now been isolated worldwide (Gharehbolagh et al. 2020), but mostly from African women either living in Africa or having immigrating to western countries (Fakhim et al. 2020). Since many reports do not mention the origin of the patients, a definitive conclusion on the spread of *C. africana* out of the African continent cannot be drawn. French Guiana is a French overseas department, located on the northeast coast of the South American continent. The population is multi-ethnic, with Amerindians, Creoles (Guyanese and Antilleans), Metropolitans and the descendants of African slaves who escaped from the plantations of Dutch Guiana in the 18th and 19th centuries (Epelboin et al. 2016).

In this study, we first aimed to describe the epidemiology of *C. africana* in vaginal specimens sampled from Guyanese women. Current methods used for the identification of yeast species, including mass spectrometry MALDI-TOF, do not allow the specific identification of *C. africana* (Hazirolan et al. 2017). Therefore, our first objective was done thanks to fluorescentintercalating-dye-real time PCR we specifically set-up. In a second instance, *C. africana* strains were subjected to molecular typing to analyze their genetic diversity and relatedness to previously characterized *C. africana* strains collected in different continents.

## Material and methods

### Collection of samples, strains and patient’s data

This study was conducted from May to August 2019 at the Centre Hospitalier Andrée Rosemont, Cayenne, French Guiana. All vaginal swabs received at the lab during this period were included in this study. Swabs were subjected to microscopic examination after gram staining and inoculated onto ChromAgar plates (Becton Dickinson, France) for a 2-day-incubation time at 35°C. Colonies had their identification confirmed thanks to Matrix-Assisted Laser Desorption/Ionisation-Time of Flight mass spectrometry (MALDI-TOF MS) analysis (Microflex Bruker). In the case of positive culture for *C. albicans*, clinical and epidemiologic data were retrospectively collected: patient demographics, clinical manifestations, <u>diabetes mellitus, pregnancy,</u> and HIV serological status. Based on clinical data, a classification was made into (i) probable vaginal candidiasis when one symptom or more suggestive of vaginal candidiasis were observed (vaginal discharge and/or itching and/or burning and/or vaginal or vulvar irritation), and (ii) probable vaginal colonization, when no vaginal symptoms were mentioned (**Figure 1**). Isolates were stored in cryovials (Microbank, ProLab Diagnostics) at -80°C before further analysis.

**Figure 1:**
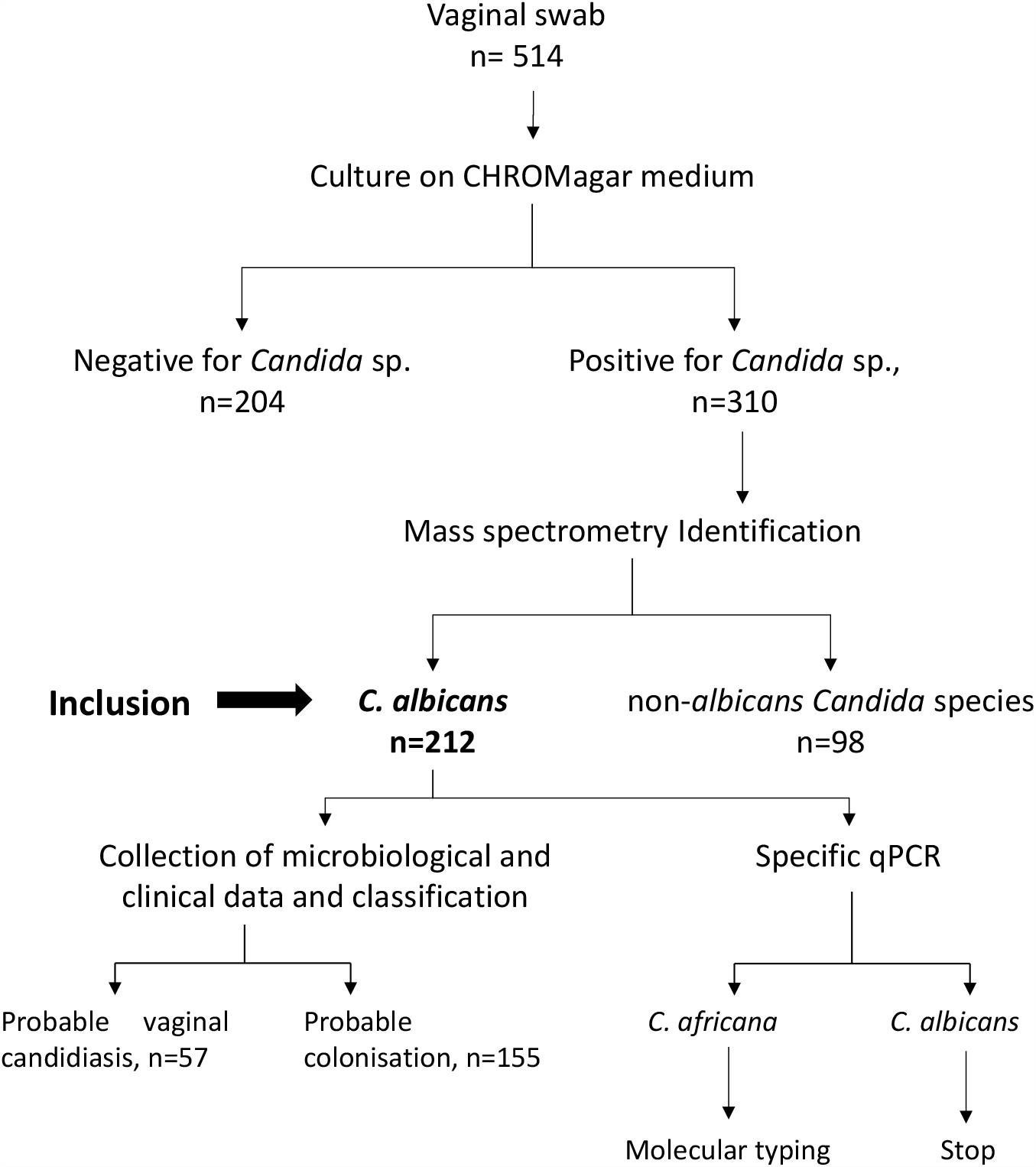
Flow chart of the study. To be classified in probable vaginal candidiasis group, the presence of one of the following suggestive symptoms is sufficient: vaginal discharge and/or Itching and/or burning and/or vaginal or vulvar irritation. Otherwise the women were classified in probable colonization group.

### Molecular biology identification

Hyphal wall protein 1 (*hwp1*) gene is highly induced during germ tube formation. A molecular method has been described to distinguish between *C. albicans* and *C. africana* based on their respective *hwp1* fragment size (Romeo et Criseo 2008). We took advantage of this trait to set up a molecular approach based on the determination of the denaturation temperature of a genomic fragment within the *hwp1* gene to distinguish *C. africana* from *C. albicans* strains belonging to other *C. albicans* clades. After a 2-day incubation at 35°C on ChromAgar plates, DNA was extracted using a rapid method with Chelex resin and heat shock (Hennequin et al. 1999). A primer pair targeting a 88-175 bp fragment within the *hwp1* gene was designed (hwp1_hrmF 5’-3’ TCATCAGCTCCTGCCACTGAA and hwp1_hrmR 5’-3’ CCGGAATAGTAATAGCACCACTT) and used in a qPCR using a ready-to-use mix containing a fluorescent-intercalating dye (EvaGreen® Fast qPCR Master Mix, Biotium, San Francisco, USA). After the amplification steps, a heat-induced dissociation of the amplicon followed by renaturation of double-strand DNA was performed allowing the determination of a melting temperature (Tm). The amplification was performed with 1X EvaGreen mix and 0.8 µM of each primer. PCR conditions were as follows: 40 cycles of denaturation at 95°C for 20 s, primer annealing and extension at 58°C for 30 s followed by dissociation step in a thermal cycler (Stratagene Mx3005P). Finally, a dissociation (melting) curve was obtained thanks to a thermal cycle consisting in a step of 95°C for 1 min, followed by 58°C for 30 s with gradual increment from 58°C to 95°C with 2 degrees per min and 95°C for 30 sec.

### Multilocus sequence typing

Strains identified as *C. africana* were then subjected to a multilocus sequence typing (MLST) protocol previously published (Bougnoux et al. 2003). This includes seven loci corresponding to housekeeping genes, namely: AAT1a, ACC1, ADP1, MPIb, SYA1, VPS13, and ZWF1b. Amplification was performed in a final volume of 25 µL containing 1X buffer, 0.3 U of Taq DNA polymerase (Qiagen), 0.25 mM deoxynucleoside triphosphate mix, 0.8 µM of each primer and 2 µL of DNA. A similar protocol of amplification was used for all targets on a Thermal cycler: denaturation for 2 min at 94°C, followed by 35 cycles of denaturation at 94°C for 1 min, annealing at 52°C for 1 min, elongation at 72°C for 1 min and a final extension step of 10 min at 72°C. Amplification products were then subjected to direct sequencing using a BigDye Terminator V3.1 kit (Life Technologies, Saint Aubin, France) with the same primers and run in a 3500xL Dx genetic analyzer (Life Technologies). The seven loci were sequenced both strands and chromatograms were manually edited and aligned using BioEdit software version 7.0.9.0. Allele numbers for each locus were assigned based on comparison of sequences obtained in this study to those deposited at the public MLST database (http://pubmlst.org/calbicans/). The composite profile of all seven allele numbers for an individual isolate defined the isolate’s diploid sequence type (DST). Fasta sequences from this study and others recovered from the mlst.net website were then used to build a Neigbhor joining tree (MEGA11.0.13).

### Ethics

For this study, an authorization has been made to the clinical investigation center and filed on June 14th 2019 (CIC, Antilles Guyane, Pr Mathieu Nacher). No ethnic information was recorded. All data were treated after anonymization.

### Data availability

https://pubmlst.org/bigsdb?db=pubmlst_calbicans_seqdef&page=alleleInfo&locus=MPIb&allele_id=191

## Results

### Clinical characteristics of patients

From May to August 2019, 514 vaginal swabs were received with 310 (60%) being culture positive for *Candida* spp (**Figure 1**). A total of 212 (68%) isolates collected from 176 women were identified as *C. albicans*. Epidemiological characteristics of those patients are shown on **Table 1**. The mean age of the patients was 28 years (range: 1–92). Among them, 147, 9 and 24 patients were pregnant, seropositive for the HIV and suffered from diabetes, respectively. Seventy-three of the 212 (34%) swabs were performed in symptomatic women suffering vulvovaginal itching (n=49, 22%), and/or burning, and/or vulvovaginal irritation (n=24, 11%). Based on these results, 57 episodes were classified in probable vaginal candidiasis, whereas the other 155 episodes were classified in probable colonization (**Table 1**).

**Table 1:**
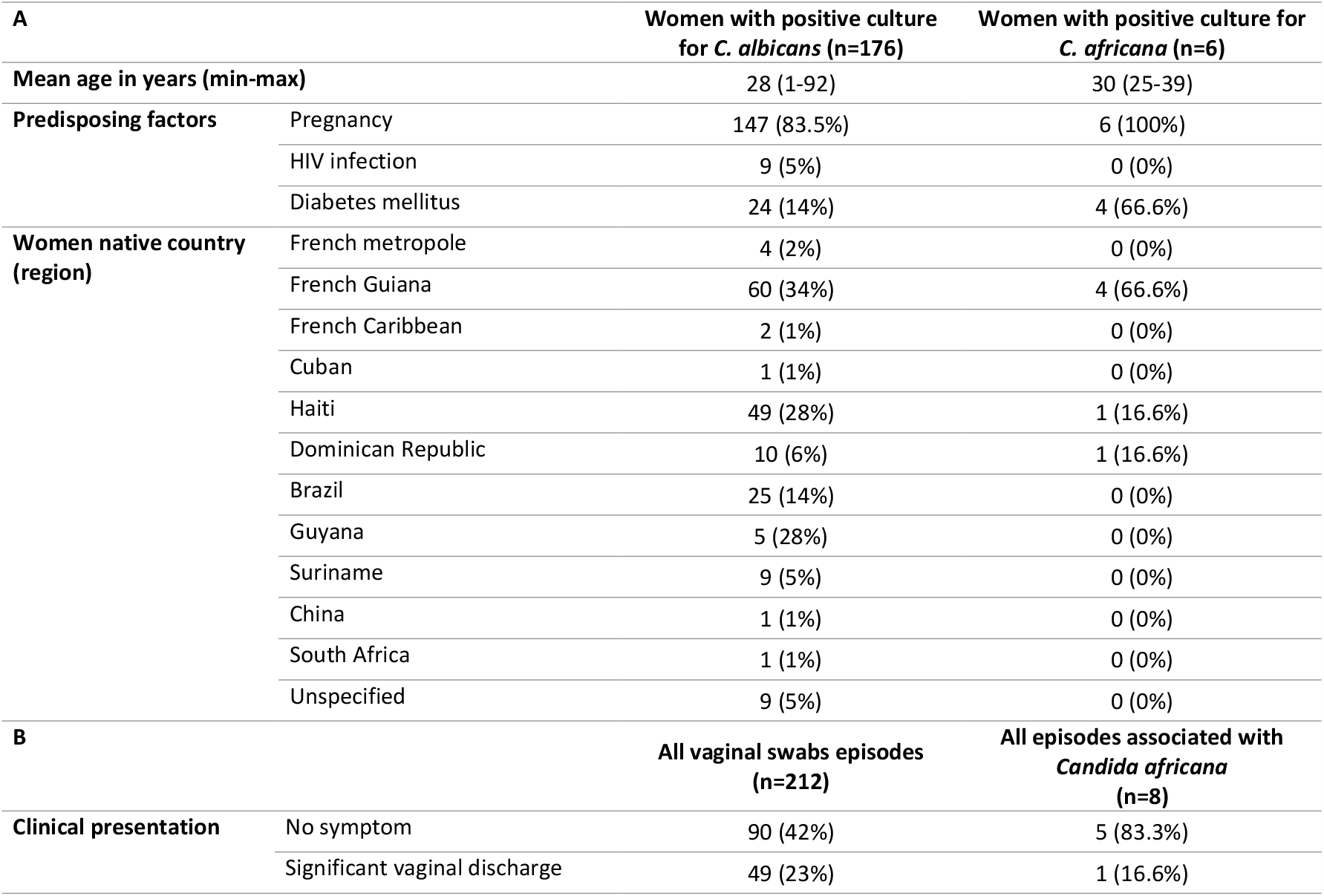
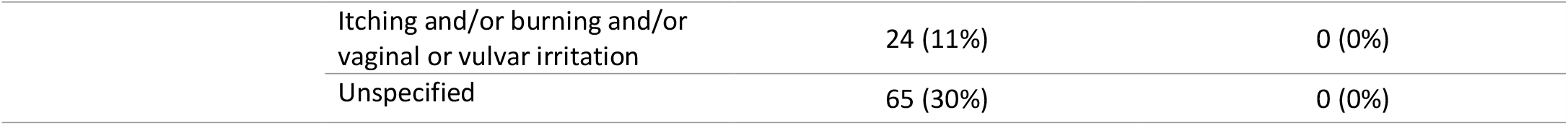
(A) Epidemiological and clinical characteristics of women who received vaginal swabs, (B) Clinical presentation associated. One woman can present different symptoms.

### Identification of Candida africana by PCR

To identify *C. africana* strains, we set up a molecular approach based on the determination of the denaturation temperature of a genomic fragment encoding the *hwp1* gene. In preliminary experiments, using reference strains, we showed that dissociation temperature of amplicons differed between *C. africana* and *C. albicans* being at 82°C and 84°C, respectively (**Figure 2**). Among the 212 *C. albicans* isolates subjected to our PCR protocol, 204 (96%) exhibited a melting temperature at 83.6 ± 0.6°C, while 8 (4%) had a melting temperature at 81.8 ± 0.2°C supporting their identification as *C. africana* (isolates G241, G355, G414, G417, G437, G468, G519 and G537). These 8 isolates had been collected from six women (3.4% of the 176 tested women). Two of them showed persistent colonization by *C. africana*, isolated twice in each case, 1 and 3 months apart, respectively. Epidemiological characteristics of those six women and clinical data corresponding to the eight episodes are presented in **Table 1**. Considering the low number of *C. africana* cases, no statistical analysis was performed. Nevertheless, we noticed that all six women were pregnant at the time of the sample, two-thirds suffered gestational diabetes and 4 were native of French Guiana while the 2 others were originally from Haiti or the Dominican Republic.

**Figure 2:**
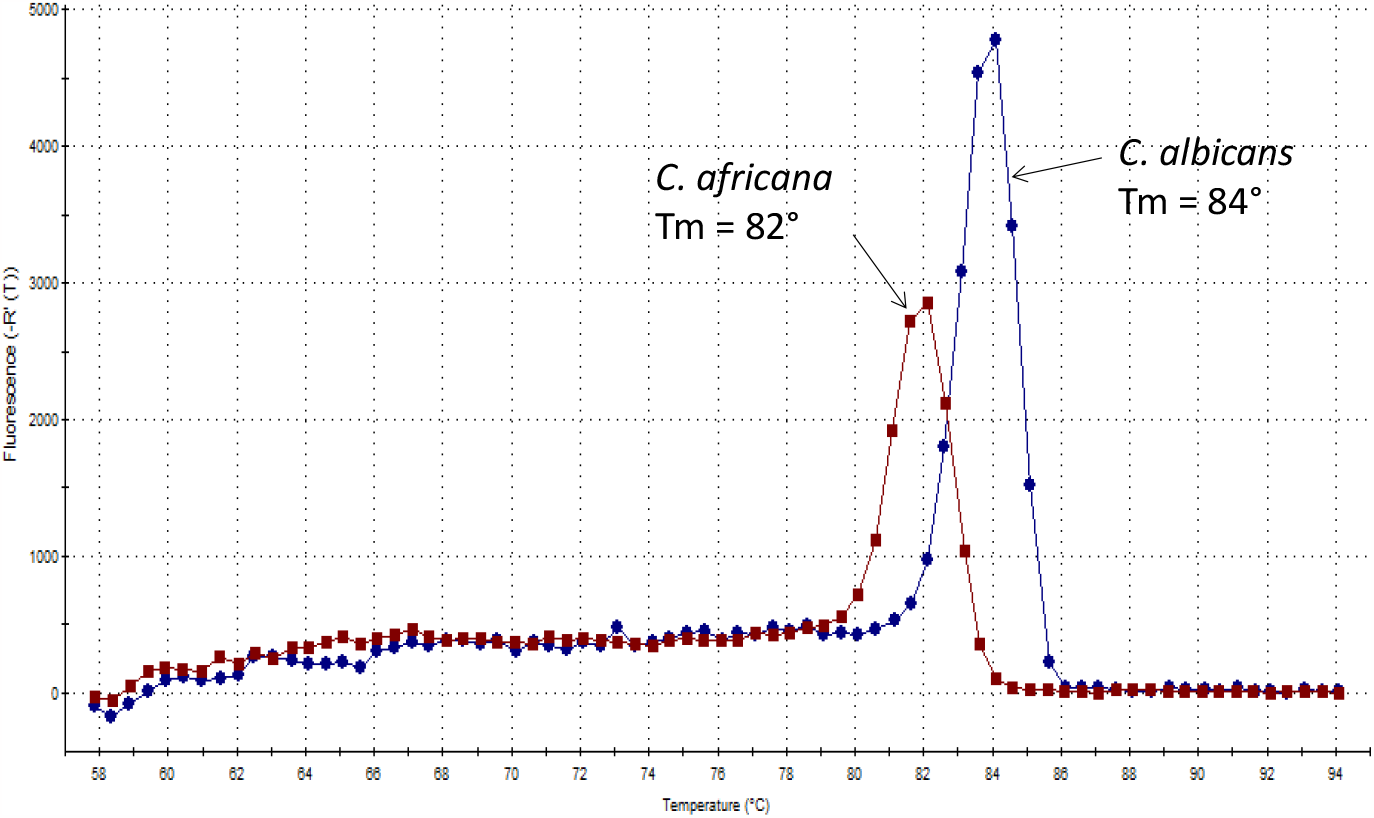
Dissociation curves of two references strains of *Candida africana* and of *Candida albicans* (ATCC90028)

### Molecular typing of Candida africana in French Guiana

The 8 isolates identified as *C. africana* were subjected to MLST. Their molecular pattern confirmed that these isolates belonged to the clade 13 of *C. albicans*. Alleles of the seven loci and their corresponding DST are summarized in **Table 2**. All alleles retrieved in this study have already been reported in the pubmslst.org database, except one of MPIb (sequence deposited in pubmlst under the number BIGSdb_20231024164852_3432492_48293). The MLST analysis revealed DST182 as the predominant genotype (5 out of 6 non-duplicated isolates). G355 and G437, two isolates collected from a single woman distantly from 3 months, had the same new DST. Concatenated sequences of the 7 loci from our study and retrieved from the pubmlst.org were used to build a neighbor-joining phylogenetic tree. Representative sequences from clade 1, 2, 3, 4 and 11 of *C. albicans* were also included. Topology of the phylogenetic tree confirmed the clustering of our strains within the clade 13 with G355 and G437 isolates being branched early in the tree (**Figure 3**).

**Table 2:** Allele number for each locus and DST, for each isolate of *Candida africana* identified during this study. DST: diploid sequence type, NDST: new DST. Isolates with * or # have been isolated from the same women.

| Isolate | DST | AAT1 | ACC1 | ADP1 | MPIb | SYA1 | VPS13 | ZWF1b |
| --- | --- | --- | --- | --- | --- | --- | --- | --- |
| **G241** | 182 | 33 | 7 | 32 | 26 | 2 | 61 | 48 |
| **G355\*** | NDST | 33 | 7 | 32 | New | 2 | 61 | 48 |
| **G414** | 182 | 33 | 7 | 32 | 26 | 2 | 61 | 48 |
| **G417#** | 182 | 33 | 7 | 32 | 26 | 2 | 61 | 48 |
| **G437\*** | NDST | 33 | 7 | 32 | New | 2 | 61 | 48 |
| **G468** | 182 | 33 | 7 | 32 | 26 | 2 | 61 | 48 |
| **G519#** | 182 | 33 | 7 | 32 | 26 | 2 | 61 | 48 |
| **G537** | 182 | 33 | 7 | 32 | 26 | 2 | 61 | 48 |

**Figure 3:**
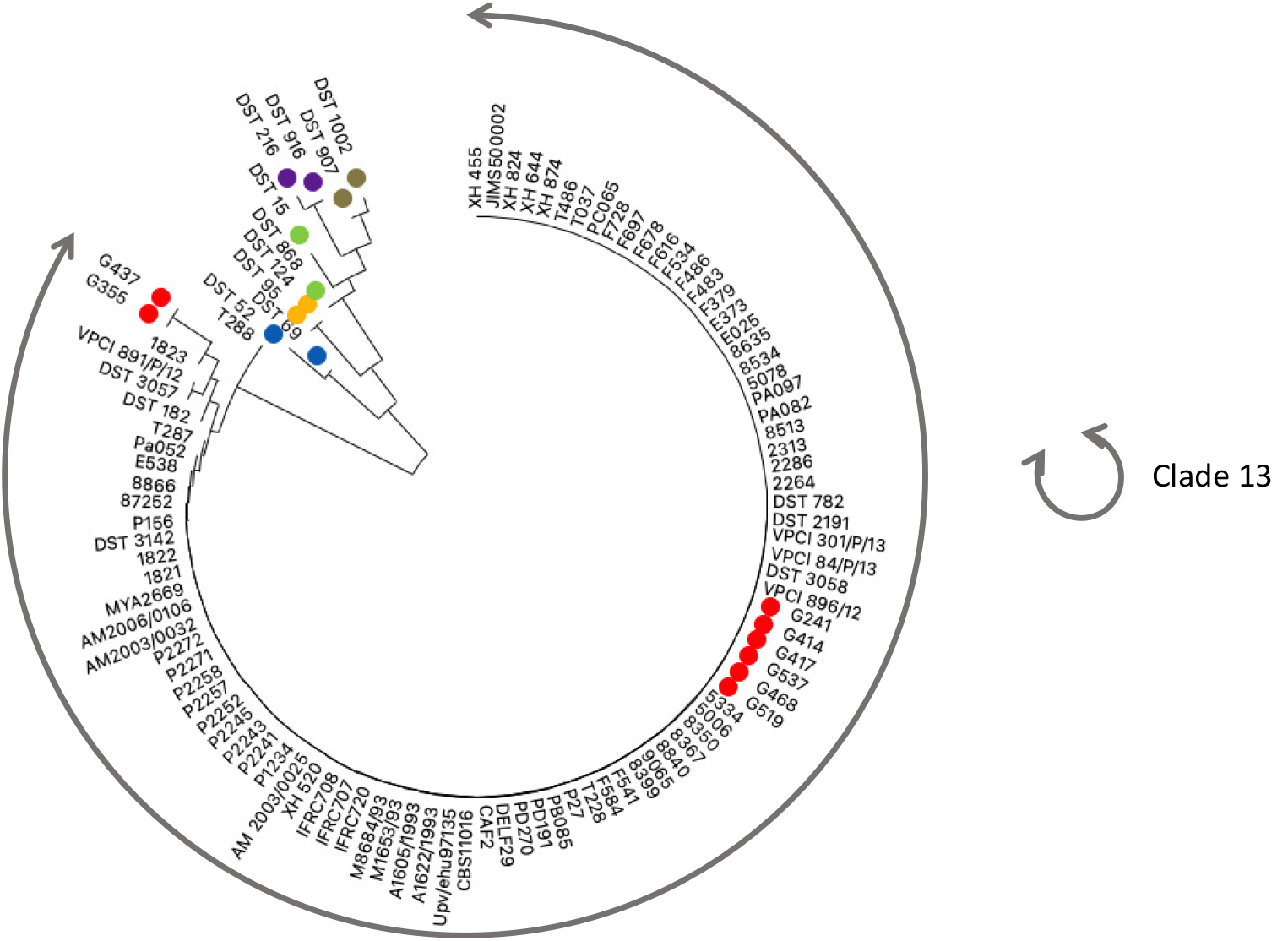
Neighbor-joining tree built using 95 *C. africana* concatenated multilocus sequences (diploid sequence type) obtained from Zhu et al 2019, PubMLST.org (clade 13) and sequences from this study (red dot). In addition, DST from clade 1 (blue dot), 2 (violet dot), 3 (green dot), 4 (yellow dot) and 11 (brown dot) were added. Phylogenetic relationship was inferred using the Neighbor-Joining method with the optimal tree shown. Distances were computed using the Maximum Composite Likelihood method using in MEGA11.0.13.

## Discussion

*C. africana* has been shown to correspond to a specific clade, clade 13, within the *C. albicans* species. Nevertheless, genomic and epidemiologic characteristics have been reported for strains belonging to this clade. The true frequency of clade 13 is various clinical context is incompletely known due to difficulty in identification.

In our study, we wanted first to assess the prevalence of *C. africana* in women living in French Guiana. However, as *C. africana* cannot be routinely identified but by using molecular approach, we therefore first developed a method based on the determination of a specific melting temperature, obtained after amplification of a *hwp1* gene fragment. This simple method can be easily implemented in labs looking at detecting *C. africana* either in collections of *C. albicans* strains or prospectively. It appears reliable as all the strains identified as *C. africana* were further confirmed to belong to the clade 13 of *C. albicans* based on MLST results. Of note, we applied the protocol directly on colonies and obtained similar results (data not shown). DST182 was the predominant genotype in our work in accordance to all previous studies conducted worldwide (Chowdhary et al. 2017). This suggests a clonal expansion, possibly through immigration.

*C. africana* infection has been primarily reported as vaginitis during the 90’s in Africa (Angola, Madagascar) (Tietz et al. 1995; Forche et al. 1999). Since 2000s, a growing number of studies reported the identification of *C. africana* worldwide. Gharehbolagh *et al*. reviewed the prevalence status of *C. africana* confirmed by molecular identification between 2001 and 2020. Cases came from 11 different countries and 4 continents with isolation from the vaginal tract in 92.8% of the cases (Gharehbolagh et al. 2020). In the different clinical settings, the prevalence of *C. africana* ranged between 0% (Malaysia) to 8,4% (Iran) (Yazdanpanah et Khaithir 2013; Naeimi et al. 2018).

In our survey, the prevalence of *C. africana* among women presenting *C. albicans* in vaginal samples was calculated at 3.4%, the highest rate observed in South America to date. This contrast with 2 previous studies conducted in South America, looking at the epidemiology of *C. africana* on vaginal samples, performed in Argentina. The first failed to detect any *C. africana* strain in a cohort of 52 pregnant women (Mucci et al. 2017), whereas the second retrieved only one *C. africana* strain among 287 (0.3%) strains obtained from patients having vaginal infection symptoms (Theill et al. 2016). While ethnicity was not recorded in our study, it should be noted that the Bushinengue population, originating from black slaves initially installed in Surinam, may represent today 30% of the Guyanese population (Price 2018). Interestingly, the new DST, described in our study, was isolated twice from a patient born in the Dominican Republic. To the best of our knowledge, there is no study focused on the epidemiology of *C. africana* in the Caribbean where the population of African origin is heavily represented.

Similar to our own survey, data from studies conducted out of African countries do not include information regarding the ethnicity of the patients (Borman, Iran, Chine). Whether virulence traits of *C. africana* could explain its propensity to colonize the vaginal tract or its inefficiency to provoke other types of infection, and if a link between ethnicity and this particular clade exists, could be further explored. Results of these studies could lead to a better understanding of host-*Candida africana* interactions.

## References

Borman, Andrew M., Adrien Szekely, Chistopher J. Linton, Michael D. Palmer, Phillipa Brown, et Elizabeth M. Johnson. 2013. « Epidemiology, Antifungal Susceptibility, and Pathogenicity of Candida Africana Isolates from the United Kingdom ». Journal of Clinical Microbiology 51 (3): 967–72. 10.1128/JCM.02816-12.

Bougnoux, M.-E., A. Tavanti, C. Bouchier, N. A. R. Gow, A. Magnier, A. D. Davidson, M. C. J. Maiden, C. d’Enfert, et F. C. Odds. 2003. « Collaborative Consensus for Optimized Multilocus Sequence Typing of Candida Albicans ». Journal of Clinical Microbiology 41 (11): 5265–66. 10.1128/JCM.41.11.5265-5266.2003.

Chowdhary, Anuradha, Ferry Hagen, Cheshta Sharma, Abdullah M. S. Al-Hatmi, Letterio Giuffrè, Domenico Giosa, Shangrong Fan, et al. 2017. « Whole Genome-Based Amplified Fragment Length Polymorphism Analysis Reveals Genetic Diversity in Candida africana ». Frontiers in Microbiology 8 (avril): 556. 10.3389/fmicb.2017.00556.

Denning, David W, Matthew Kneale, Jack D Sobel, et Riina Rautemaa-Richardson. 2018. « Global Burden of Recurrent Vulvovaginal Candidiasis: A Systematic Review ». The Lancet Infectious Diseases 18 (11): e339–47. 10.1016/S1473-3099(18)30103-8.

Epelboin, Loïc, Tomasz Chroboczek, Emilie Mosnier, Philippe Abboud, Antoine Adenis, Denis Blanchet, Magalie Demar, et al. 2016. « L’infectiologie en Guyane : le dernier bastion de la médecine tropicale française ». La Lettre de l’Infectiologue Tome XXXI-n° 4-juillet-août 2016. http://www.hal.inserm.fr/inserm-01407150/document.

Fakhim, H., A. Vaezi, J. Javidnia, E. Nasri, D. Mahdi, K. Diba, et H. Badali. 2020. « Candida Africana Vulvovaginitis: Prevalence and Geographical Distribution ». Journal de Mycologie Médicale 30 (3): 100966. 10.1016/j.mycmed.2020.100966.

Felice, Maria Rosa, Megha Gulati, Letterio Giuffrè, Domenico Giosa, Luca Marco Di Bella, Giuseppe Criseo, Clarissa J. Nobile, Orazio Romeo, et Fabio Scordino. 2016. « Molecular Characterization of the N-Acetylglucosamine Catabolic Genes in Candida africana, a Natural N-Acetylglucosamine Kinase (HXK1) Mutant ». PLoS ONE 11 (1): e0147902. 10.1371/journal.pone.0147902.

Forche, Anja, Gabriele Schönian, Yvonne Gräser, Rytas Vilgalys, et Thomas G. Mitchell. 1999. « Genetic Structure of Typical and Atypical Populations of Candida Albicans from Africa ». Fungal Genetics and Biology 28 (2): 107–25. 10.1006/fgbi.1999.1164.

Gharehbolagh, Sanaz Aghaei, Bahareh Fallah, Alireza Izadi, Zeinab Sadeghi Ardestani, Pooneh Malekifar, Andrew M. Borman, et Shahram Mahmoudi. 2020. « Distribution, Antifungal Susceptibility Pattern and Intra-Candida Albicans Species Complex Prevalence of Candida Africana: A Systematic Review and Meta-Analysis ». Édité par Joy Sturtevant. PLOS ONE 15 (8): e0237046. 10.1371/journal.pone.0237046.

Hazirolan, G., H.U. Altun, R. Gumral, N.C. Gursoy, B. Otlu, et B. Sancak. 2017. « Prevalence of Candida Africana and Candida Dubliniensis, in Vulvovaginal Candidiasis: First Turkish Candida Africana Isolates from Vulvovaginal Candidiasis ». Journal de Mycologie Médicale 27 (3): 376–81. 10.1016/j.mycmed.2017.04.106.

Hennequin, C., E. Abachin, F. Symoens, V. Lavarde, G. Reboux, N. Nolard, et P. Berche. 1999. « Identification of Fusarium Species Involved in Human Infections by 28S rRNA Gene Sequencing ». Journal of Clinical Microbiology 37 (11): 3586–89.

Mucci, María Josefina, María Luján Cuestas, María Fernanda Landanburu, et María Teresa Mujica. 2017. « Prevalence of Candida Albicans, Candida Dubliniensis and Candida Africana in Pregnant Women Suffering from Vulvovaginal Candidiasis in Argentina ». Revista Iberoamericana de Micología 34 (2): 72–76. 10.1016/j.riam.2016.09.001.

Naeimi, Behrouz, Hossein Mirhendi, Gholamreza Khamisipour, Farzaneh Sadeghzadeh, et Bahram Ahmadi. 2018. « Candida Africana in Recurrent Vulvovaginal Candidiasis (RVVC) Patients: Frequency and Phenotypic and Genotypic Characteristics ». Journal of Medical Microbiology 67 (11): 1601–7. 10.1099/jmm.0.000834.

Price, Richard. 2018. « Maroons in Guyane: Getting the Numbers Right ». New West Indian Guide / Nieuwe West-Indische Gids 92 (3-4): 275–83. 10.1163/22134360-09203001.

Romeo, Orazio, et Giuseppe Criseo. 2008. « First molecular method for discriminating between Candida africana, Candida albicans, and Candida dubliniensis by using hwp1 gene ». Diagnostic Microbiology and Infectious Disease 62 (2): 230–33. 10.1016/j.diagmicrobio.2008.05.014.

Sharma, Cheshta, Sumathi Muralidhar, Jianping Xu, Jacques F. Meis, et Anuradha Chowdhary. 2014. « Multilocus Sequence Typing of Candida Africana from Patients with Vulvovaginal Candidiasis in New Delhi, India ». Mycoses 57 (9): 544–52. 10.1111/myc.12193.

Theill, Laura, Catiana Dudiuk, Susana Morano, Soledad Gamarra, María Elena Nardin, Emilce Méndez, et Guillermo Garcia-Effron. 2016. « Prevalence and Antifungal Susceptibility of Candida Albicans and Its Related Species Candida Dubliniensis and Candida Africana Isolated from Vulvovaginal Samples in a Hospital of Argentina ». Revista Argentina de Microbiología 48 (1): 43–49. 10.1016/j.ram.2015.10.003.

Tietz, H J, A Küssner, M Thanos, M P De Andrade, W Presber, et G Schönian. 1995. « Phenotypic and genotypic characterization of unusual vaginal isolates of Candida albicans from Africa. » Journal of Clinical Microbiology 33 (9): 2462–65.

Yazdanpanah, Atta, et Tzar Mohd Nizam Khaithir. 2013. « Issues in Identifying Germ Tube Positive Yeasts by Conventional Methods ». Journal of Clinical Laboratory Analysis 28 (1): 1–9. 10.1002/jcla.21635.

